# Functional Changes Induced by Short Term Caloric Restriction in Cardiac and Skeletal Muscle Mitochondria

**DOI:** 10.1101/2020.03.13.991232

**Authors:** Julian David C. Serna, Camille C. Caldeira da Silva, Alicia J. Kowaltowski

## Abstract

Caloric restriction (CR) is widely known to increase life span and resistance against different types of injuries in several organisms. We have previously shown that mitochondria from livers or brains of CR animals exhibit higher calcium uptake rates and lower sensitivity to calcium-induced mitochondrial permeability transition (mPT), an event related to the resilient phenotype exhibited by these organs. Given the importance of calcium in metabolic control and cell homeostasis, we aimed here to uncover possible changes in mitochondrial calcium handling, redox balance and bioenergetics in cardiac and skeletal muscle mitochondria. Unexpectedly, we found that CR does not alter the susceptibility to mPT in muscle (cardiac or skeletal), nor calcium uptake rates. Despite the lack in changes in calcium transport properties, CR consistently decreased respiration in the presence of ATP synthesis in heart and soleus muscle. In heart, such changes were accompanied by a decrease in respiration in the absence of ATP synthesis, lower maximal respiratory rates and a reduced rate of hydrogen peroxide release. Hydrogen peroxide release was unaltered by CR in skeletal muscle. No changes were observed in inner membrane potentials and respiratory control ratios. Together, these results highlight the tissue-specific bioenergetic and ion transport effects induced by CR, demonstrating that resilience against calcium-induced mPT is not present in all tissues.

## Introduction

Heart and skeletal muscle exhibit several detrimental changes during aging, and are key tissues in age-related dysfunctions (McCormick *et al.*, 2018; Chiao & Rabinovitch, 2015; López-Otín *et al.*, 2013; de Melo *et al.*, 2018). Both display a high dependence on aerobic metabolism for their performance, and remarkable metabolic plasticity, with adaptations to use different substrates, as well as to changes in oxygen and nutrient supply (Lesnefski, *et al.* 2017; Chouchani *et al.*, 2014; Gillani *et al.*, 2012). Since mitochondria are central organelles in metabolic plasticity, strategies to preserve mitochondrial function in these tissues are the focus of intense research (Lázár *et.al.*, 2017).

A mitochondrial phenomenon associated with many age-related pathological conditions is the mitochondrial permeability transition (mPT), a sudden increase in inner mitochondrial membrane permeability (Hunter & Haworth, 1979a; Haworth & Hunter, 1979b, Haworth & Hunter, 1979, reviewed by Chinopoulus, 2018; Vercesi *et al.*, 2018). mPT results in loss of ion gradients and metabolite accumulation, hampering oxidative phosphorylation. Swelling and rupture of the outer mitochondrial membrane secondary to mPT may also promote cell death by necrosis or apoptosis, thus leading to tissue damage in a variety of pathological conditions (Lemasters *et al.*, 2009; Figueira *et al.*, 2013; Vercesi *et al.*, 2018).

Calcium overaccumulation in mitochondria is the central trigger for mPT, which is also stimulated by a high concentration of oxidants, protein thiol oxidation and missfolding, phosphate, and many other stimuli (Halestrap *et al.*, 2015; Figueira *et al.*, 2013; Vercesi *et al.*, 2018). mPT induction is closely related to mitochondrial oxidant release; interventions that augment antioxidant defenses and/or decrease the rate of oxidant release are expected to decrease the probability of mPT (reviewed in Kowaltowski *et al.*, 2001; Figueira *et al.*, 2013; Vercesi *et al.*, 2018). Susceptibility to mPT also increases with age in several tissues (Panel *et al.*, 2018). The same is observed with conditions that promote unhealthy aging, such as high-fat diets, obesity and diabetes (Littlejohns *et al.*, 2014; Castillo *et al.*, 2019; Anderson *et al.*, 2019). In contrast, caloric restriction (CR), a dietetic intervention that involves a decrease in energy intake without malnutrition is well known to promote an increase in lifespan and improve healthspan, and protects against calcium-induced mPT in rodent livers and brains (Kristal *et al.*, 1998; Amigo *et al.*, 2017; Menezes-Filho *et al.*, 2017). Short term CR also increases mitochondrial calcium retention capacity and uptake rates in these tissues (Amigo *et al.* 2017; Menezes *et al.* 2017). These data support the idea that mPT and mitochondrial Ca^2+^ transport are modulated by aging and diet.

Indeed, several studies demonstrate that CR decreases the rate of oxidant release by mitochondria, an important factor in mPT induction (Kowaltowski *et al.*, 2001; Figueira *et al.*, 2013; Vercesi *et al.*, 2018). However, other papers demonstrate unaltered, decreased and even increased in oxidant release with dietary restriction (Walsh *et al.* 2014), so a prevention of oxidant formation does not seem to be a consistent finding in CR. Mitochondrial bioenergetics also has been shown to be modulated by CR, preventing age-related dysfunction of the electron transport chain (ETC) in heart and different types of skeletal muscle (Hofer *et al*, 2009; Lanza *et al.*, 2012). However, calcium transport properties of heart and skeletal muscle mitochondria from CR animals have not been explored within the current framework. Here, we aimed to uncover possible alterations in mitochondrial calcium transport, mPT, bioenergetics and H_2_O_2_ release induced by CR, in heart and soleus muscle. Interestingly, we found that the effects of CR in brain and liver are not applicable to skeletal tissues, indicating that CR-induced modulation of mitochondrial Ca^2+^ transport is tissue-specific.

## Experimental Procedures

### Animal care and caloric restriction

Experiments were approved by the local committee for animal care and use and followed NIH guidelines. Male Sprague-Dawley (NTac:SD) rats were used, bred and lodged at the *Biotério de Produção e Experimentação da Faculdade de Ciências Farmacêuticas e Instituto de Química*. Animals were lodged in plastic ventilated cages (49 cm × 34 cm × 16 cm), 2 animals per cage, under specific pathogen free conditions, at constant temperature (22 ± 2°C) and relative air humidity (50 ±10%). They were exposed to a photoperiod of 12 h light / 12 h dark, with the light period beginning at 6:00. Until 12 week (adults), the animals were provided with the standard AIM93G diet (prepared by Rhoster). After that, animals were randomly divided in two groups. Following a brief period of adaptation (2 days), the daily food consumption of the CR group was established as 60% of the *ad libitum* feeding quantity. CR animals were then provided with supplemented AIM93M chow (Rhoster) to induce 40% CR without changes in vitamin and mineral levels (Cerqueira and Kowaltowski, 2010). Over the next months, food consumption was adjusted based on AL daily food consumption. Water consumption was maintained *ad libitum*.

### Euthanasia

Animals were fasted before each experiment for 10-12 h, beginning at 19:00. Before euthanasia, they were deeply anesthetized (1 mL ^.^ Kg^−1^ ketamine, 0.6 mL ^.^ Kg^−1^ xylazine and 0.5 mL ^.^ Kg^−1^ acepromazine, subcutaneously). Animals were then submitted to cardiac puncture and the diaphragm was cut. Four animals were employed per group.

### Mitochondrial isolation

Heart mitochondria were isolated as described by Gostimskaya and Galkin (2010) with a few modifications. All steps were performed at 4°C. After washing in PBS supplemented with 10 mM EDTA, connective tissues were rapidly removed. The heart was chopped into small segments and then incubated in washing buffer (0.3 M sucrose, 10 mM HEPES, 2 mM EGTA, 1 mM EDTA, pH = 7.2) plus 0.125 mg ^.^ mL^−1^ trypsin during 10 min. The mixture was homogenized in a tissue grinder (at least three strokes) and then incubated during 5 additional minutes in an additional amount of washing buffer, again supplemented with trypsin. After that, three additional strokes were performed to complete the homogenization step. The suspension was diluted with 0.2% BSA to inactivate trypsin. The mitochondrially-enriched fraction was prepared through differential centrifugation. The suspension was initially centrifuged at 600 g during 15 min, and the resultant supernatant was then centrifuged at 8500 g. The pellet was resuspended in 10 mL of isolation buffer, and again centrifuged at 8500 g for 10 minutes. The pellet was finally resuspended, including the fluffy layer (to avoid eliminating mitochondria differentially between samples), in a minimum amount of isolation buffer. Protein concentration was determined using the Bradford Method.

Skeletal muscle mitochondria were isolated, with some modifications, as previously described by Frezza *et al.* 2007). All procedures were performed at 4°C. The soleus muscle of each hind limb was collected and washed in PBS supplemented with 10 mM EDTA. After connective tissue removal, the muscles were chopped in small pieces and then incubated in muscle isolation buffer 1 (67 mM sucrose, 50 mM KCl, 1 mM EDTA, and 0.125 mg ^.^ mL^−1^ trypsin; pH = 7.4) during 10 minutes. The mixture was homogenized (3-4 strokes) and then incubated during 5 min in an additional amount of buffer, in the presence of 0.125 mg ^.^ mL^−1^ trypsin. The suspension was processed again in the tissue grinder (3 strokes) and then diluted with isolation buffer plus 0.2% BSA (fatty acid free). The suspension was centrifuged at 800 g during 15 min, and the resulting supernatant was then centrifuged at 8500 g for 10 min. The pellet was resuspended in 10 mL of isolation buffer, and centrifuged again at 8500 g. Finally, the pellet was resuspended in isolation buffer (including the fluffy layer). Protein concentration was measured through the Bradford Method.

### Ca^2+^ uptake assays

500 μg of the mitochondrially-enriched fraction (derived from heart or soleus muscle) were incubated in 2 mL of experimental buffer (125 mM sucrose, 65 mM KCl, 10 mM Hepes, 2 mM phosphate, 2 mM MgCl_2_ and 0.2% bovine serum albumin, adjusted to pH 7.2 with KOH), plus 0.1 μM Calcium Green 5N and 1 mM succinate, in the presence of 2 μM rotenone. Calcium Green 5N fluorescence was measured with a F4500 Hitachi Fluorimeter at excitation and emission wavelengths of 506 nm and 532 nm, at 37°C, under constant stirring. Several additions of 10 μM Ca^2+^, with 3 minute intervals, were made until mPT induction (Amigo *et al.*, 2017; Menezes-Filho *et al.*, 2017).

The relationship between Calcium Green 5N fluorescence (F) and [Ca^2+^] concentrations was established using the equation [Ca^2+^]=K_d_.(F-F_min_)/(F_max_-F). The K_d_ value was empirically determined as the value at which the change in fluorescence (ΔF) after each calcium addition is equivalent to a 10 μM change in calcium concentration (Δ[Ca^2+^]). The maximal (F_max_) and minimal (F_min_) fluorescence were determined at the end of each trace using 100 μL of 100 mM Ca^2+^ and 100 mM EGTA solutions, respectively. The calcium retention capacity was determined as the total amount of calcium Ca^2+^ taken up by mitochondria until the induction of mPT (nmol Ca^2+.^ mg protein^−1^). Calcium uptake rates were determined as the slope of the linear portion (nmol Ca^2+.^ mg protein^−1.^ s^−1^) at the beginning of the first calcium addition. An exponential decay fit (first order kinetics) was used to calculate the minimum calcium concentration achieved in the medium, or the affinity of mitochondria for Ca^2+^, a concentration under which calcium is not taken up by mitochondria.

### Mitochondrial oxygen consumption

Oxygen consumption was assessed using high-resolution Oroboros oxygraph. Heart mitochondria (75 μg) or 150 μg of the soleus muscle mitochondrial fraction were suspended in 2 mL of the same experimental buffer used for calcium uptake assays. Mitochondria were incubated with 1 mM succinate in the presence of 1 μM rotenone, under constant stirring at 37°C. ADP (2 mM) was used to induce State 3. State 4 and maximal respiration (state 3u, uncoupled) were induced with 1 μM oligomycin (muscle and heart mitochondria) and CCCP (1.25 μM or 2 μM for heart or soleus mitochondria), respectively. Respiratory control ratios were determined as State 3 divided by State 4.

### Inner mitochondrial membrane potential measurements

Mitochondrial membrane potentials (ΔΨ) were estimated through changes in the fluorescence (quenching mode) of the safranin O dye (Akerman & Wikström, 1976). 75 μg of heart mitochondria (or 100 μg of soleus muscle mitochondria) were incubated in 2 mL experimental buffer in the presence of 5 μM Safranin O, 1 mM succinate and 1 μM rotenone (State 2), pH = 7.4, adjusted with KOH. The ΔΨ values different respiratory states were assessed as described above. Fluorescence was followed in an F2500 Hitachi Fluorimeter at excitation and emission wavelengths of 485 nm and 586 nm, respectively, under constant stirring, at 37°C. The dye is accumulated in the mitochondrial matrix in a ΔΨ-dependent fashion; membrane depolarization leads to an increase in net fluorescence. A calibration curve was employed to establish a relationship between ΔΨ and fluorescence (Kowaltowski *et al.*, 2002). ΔΨ was clamped at different values through changes in extramitochondrial K^+^ concentrations, in an initially K^+^-free experimental media.

### Hydrogen peroxide release from isolated mitochondria

Mitochondrial H_2_O_2_ release was followed through the oxidation of Amplex Red (25 μM), a reaction catalyzed by horseradish peroxidase (5 U/mL; HRP; Zhou *et al.*, 1997). 75 μg of heart mitochondria or 100 μg soleus muscle mitochondria were added to 2 mL of experimental media in the presence of 1 mM succinate and 1 μM rotenone under constant stirring, at 37°C. The oxidation of Amplex Red generates a fluorescent compound (resorufin), which was detected with a F2500 Hitachi Fluorimeter at a excitation and emission wavelength of 563 nm and 587 nm, respectively. A calibration curve, used to establish a relationship between resorufin fluorescence and H_2_O_2_ concentration, was constructed under the same experimental conditions (in the absence of mitochondria), by sequentially adding known quantities of H_2_O_2_.

### Statistical analysis

Results are presented as means ± SEM and were analyzed using GraphPad Prism Software 4.0. Data were compared using t-tests.

## Results

### CR does not change heart mitochondrial calcium transport properties

We have previously found that CR increases Ca^2+^ uptake rates and resistance against calcium-induced mPT in the liver and brain (Amigo *et al.*, 2017; Menezes-Filho *et al.*, 2017). This lead us to investigate possible changes in heart mitochondrial calcium handling promoted by CR (**Fig. 1**). We isolated a mitochondrial fraction containing both subsarcolemmal and intermyofibrillar populations of these organelles (Palmer *et al.*, 1977). These mitochondria were incubated in the presence of Ca^2+^ Green 5N, which fluoresces upon calcium binding but is not membrane-permeable, thus allowing us to detect changes in the calcium concentrations of the extramitochondrial compartment. **Fig. 1A** shows a typical calcium uptake trace of mitochondria obtained from hearts of *ad libitum* (AL, blue trace) or CR (red trace) rats. After each 10 μM addition of calcium, marked by an arrow, and acute increases in [Ca^2+^] promoted by the addition, a gradual decrease in extra-mitochondrial calcium concentration is observed, indicative of uptake of the ion by mitochondria. Each trace finished after induction of mPT, marked by a sudden increase in extramitochondrial calcium (release of stored calcium). No differences in Ca^2+^ retention capacity were noted between the AL and CR groups (**Fig. 1B** shows quantified comparisons).

**Figure 1.**
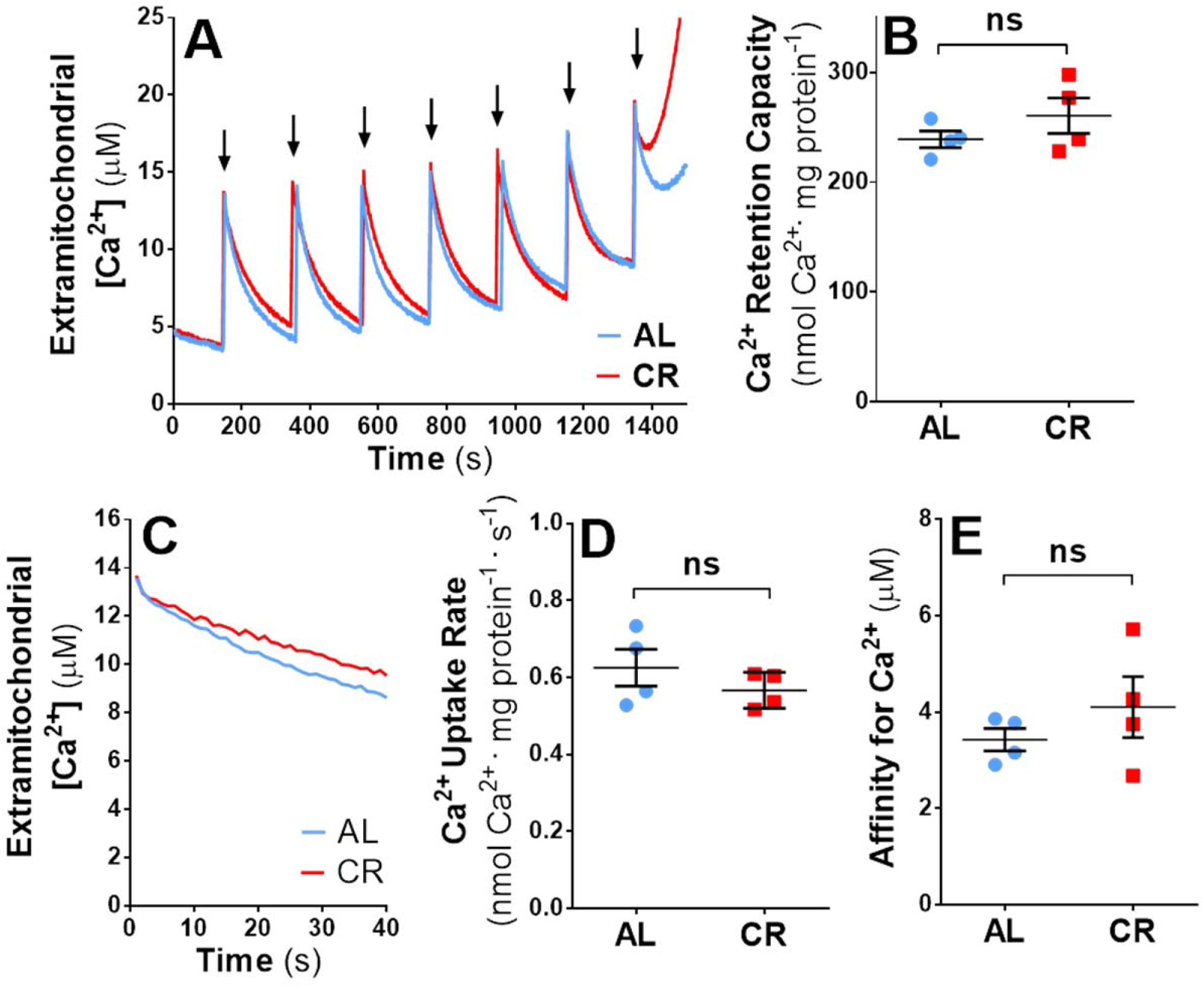
CR does not change heart mitochondrial Ca^2+^ retention capacity and uptake rate. 500 μg of total mitochondrial protein were incubated in 2 mL media, as described in the methodology. **A.** Representative Ca^2+^ uptake traces of samples derived from AL (blue) or CR (red) animals; each arrow represents a 10 μM calcium addition. **B.** Quantification of maximal Ca^2+^ retention capacity derived from graphs such as shown in Panel A. **C.** Representative mitochondrial Ca^2+^ uptake trace after the first Ca^2+^ addition. **D.** Quantification of initial mitochondrial Ca^2+^ uptake rates derived from plots such as shown in panel C. **E.** Data such as from Panel A were fitted with a first order exponential decay to calculate the minimum [Ca^2+^] reached in the medium, or the affinity of these mitochondria for Ca^2+^. ns = non-significant.

We also investigated possible changes in the kinetics of calcium clearance by mitochondria. **Fig. 1C** shows an initial calcium uptake rate trace, quantified in **Fig. 1D**. Under the experimental conditions employed, we were not able to detect any diet-induced changes in the calcium uptake rate. Finally, the affinity of mitochondria for Ca^2+^ ([Ca^2+^] in the media after uptake stabilized) was equal in CR and AL groups (**Fig. 1E**). Overall, heart mitochondrial Ca^2+^ uptake properties and their modulation by CR differ significantly in heart from brain or liver (Amigo *et al.*, 2017; Menezes-Filho *et al.*, 2017), suggesting that the effects of CR on mitochondrial Ca^2+^ uptake are tissue-specific.

### CR alters mitochondrial oxygen consumption and redox state without changes in coupling or the inner membrane potential

The ability of CR to preserve or increase bioenergetic fitness in heart mitochondria over time is debated (Walsh *et al.* 2014). We investigated possible changes in mitochondrial oxygen consumption using an Oroboros high resolution respirometer (**Fig. 2**). When ADP was added to induce oxidative phosphorylation (state 3 respiration), CR mitochondria displayed lower respiratory rates (**Fig. 2A**). The same was observed when ATP synthase was inhibited by oligomycin (state 4 respiration) or maximized by the uncoupler CCCP (maximal respiration), demonstrating that CR limits electron transport rates in heart mitochondria. Interestingly, these changes were not accompanied by any alteration in respiratory control ratios (**Fig. 2B**), which determine coupling between respiration and ATP synthesis, measured as State 3/State 4. Indeed, calibrated inner mitochondrial membrane potentials (ΔΨ, Akerman, 1976; Kowaltwski *et al.*, 2002) were identical in AL and CR mitochondria (**Fig. 2C**). Since ΔΨ is the driving force for Ca^2+^ accumulation, this lack of change may explain the equal Ca^2+^ uptake rates observed in **Fig. 1**.

**Figure 2.**
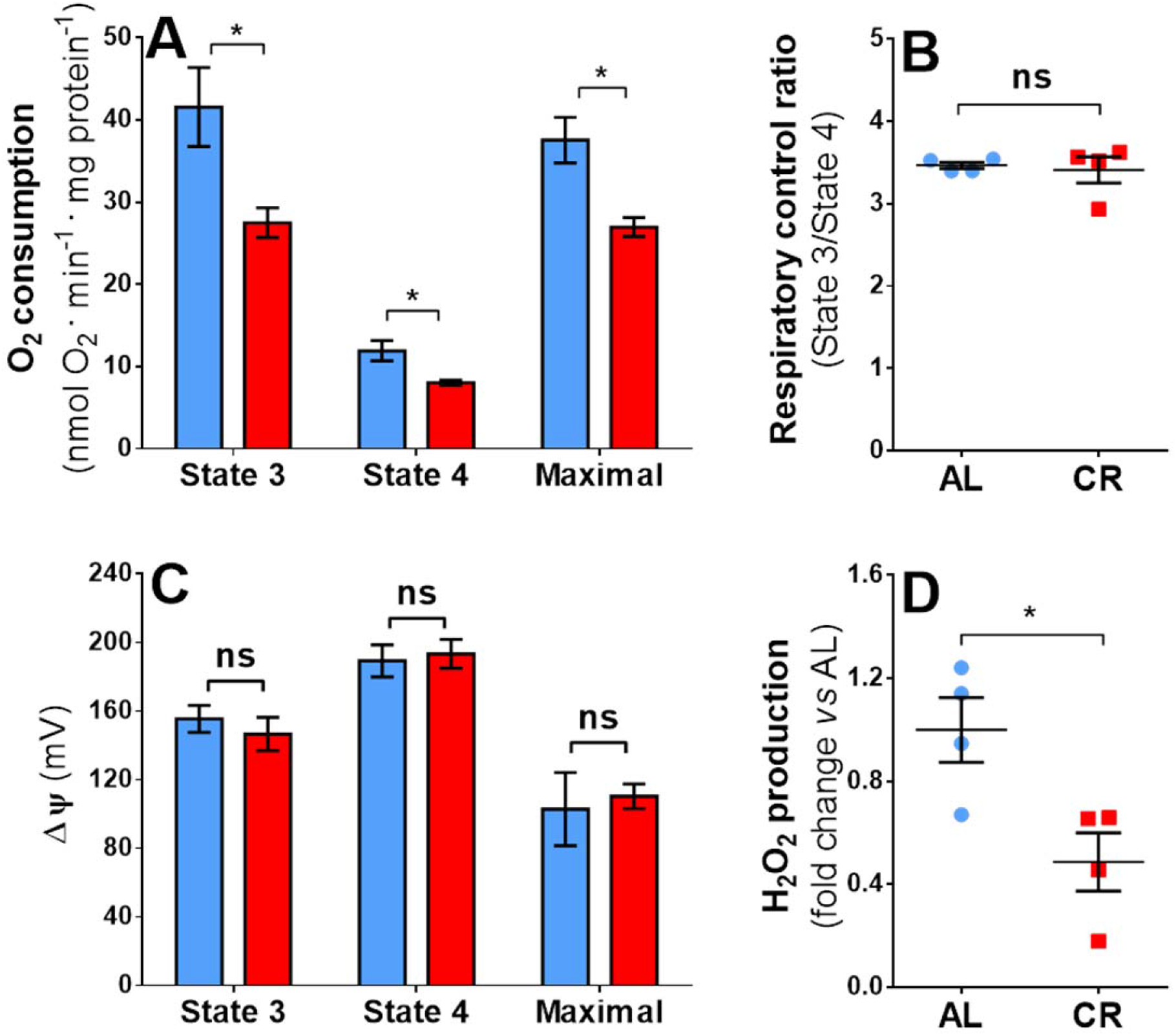
CR modulates heart mitochondrial oxygen consumption and H_2_O_2_ release without changes in coupling or inner membrane potentials. Mitochondrial oxygen consumption, membrane potentials (ΔΨ) and peroxide release were measured as described in the methodology. Blue or red bars and points represent data obtained from AL or CR animals, respectively. **A.** Oxygen consumption rates were measured in mitochondria incubated, successively, in the presence of succinate and rotenone plus ADP (State 3), oligomycin (State 4) and CCCP (Maximal). **B.** Respiratory control ratios (State 3/State 4). **C.** Membrane potentials, calibrated in mV. **D.** Hydrogen peroxide release normalized to average AL fluorescence. *p<0.05 versus AL. ns = non-significant, * = p < 0.05.

Changes in oxygen consumption or ΔΨ are often associated with altered mitochondrial oxidant release (Korshunov *et al.*, 1997; Caldeira da Silva *et al.* 2008; Tahara et al., 2009). We consequently measured possible changes in H_2_O_2_ release by mitochondria using Amplex Red and horseradish peroxidase (Zhou *et al.*, 1997). CR promoted a substantial decrease in H_2_O_2_ release by heart mitochondria (**Fig. 2D**).

### CR does not change soleus muscle mitochondrial calcium transport properties

We tested the ability of CR to promote changes in calcium handling in skeletal muscle mitochondria. Mitochondria were isolated from the soleus muscle, enriched in type I fibers and with high dependence on aerobic metabolism (Soukup *et al.*, 2002), using a preparation containing intermyofibrillar and subsarcolemmal populations (Kayar *et al.*, 1988). **Fig. 3A** depicts a representative calcium uptake trace, quantified in **Fig. 3B.** A remarkably higher (around 50%) calcium retention capacity was seen in soleus mitochondria when compared to heart mitochondria, but CR did not promote changes in maximal Ca^2+^ uptake and the sensibility to calcium-induced mPT. Calcium uptake rates in soleus mitochondria (**Fig. 3C, 3D**) were also unchanged by CR, but around 70% higher than for heart mitochondria. Finally, the affinity for Ca^2+^ was unchanged by CR, but higher in skeletal muscle than in heart (**Fig. 3E**).

**Figure 3.**
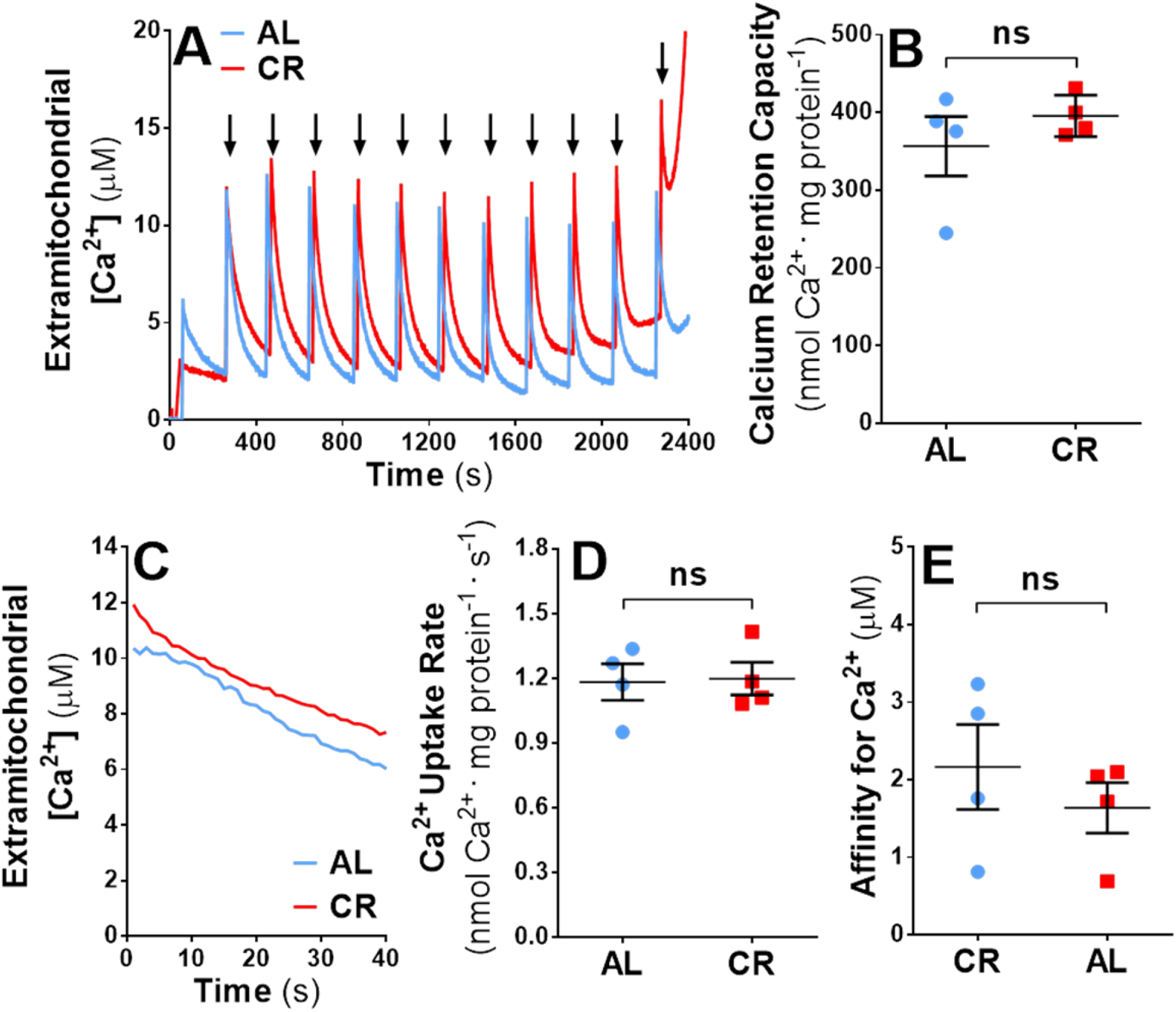
CR does not change soleus muscle mitochondrial Ca^2+^ retention capacity and uptake rate. 500 μg of total mitochondrial protein were incubated in 2 mL media, as described in the *Experimental* section. **A.** Representative Ca^2+^ uptake traces of samples derived from AL (blue) or CR (red) animals; each arrow represents a 10 μM calcium addition. **B.** Quantification of maximal Ca^2+^ retention capacity derived from graphs such as shown in Panel A. **C.** Representative mitochondrial Ca^2+^ uptake traces after the first Ca^2+^ addition. **D.** Quantification of initial mitochondrial Ca^2+^ uptake rates derived from plots such as shown in panel C. **E.** Data such as from Panel A were fitted with a first order exponential decay to calculate the minimum [Ca^2+^] achieved in the medium, or the affinity of these mitochondria for Ca^2+^. ns = non-significant.

### CR decreases mitochondrial state 3 oxygen consumption without changes in coupling, membrane potentials or H_2_O_2_

As previously observed in heart mitochondria, CR promotes a decrease in skeletal muscle respiration under state 3 conditions (**Fig. 4A**). No changes in state 4 and maximal respiration were observed as a result of the CR diet. Mitochondrial respiratory control ratios in muscle were also unchanged as result of the dietary intervention, **Fig. 4B**, as were ΔΨ and H_2_O_2_ release, **Fig. 4C** and **4D**.

**Figure 4.**
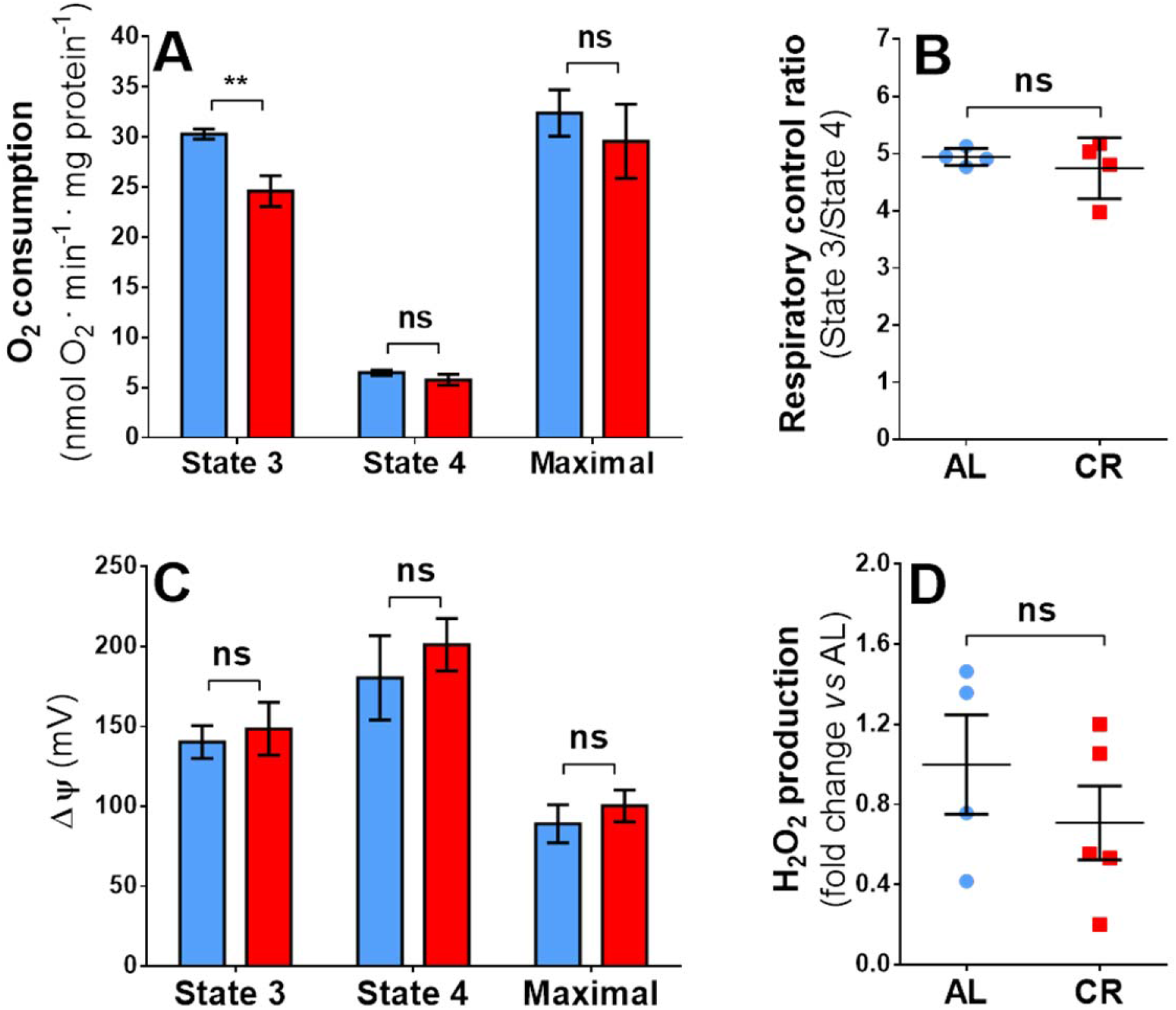
CR alters state 3 mitochondrial oxygen consumption without changes in coupling, membrane potentials (ΔΨ) or H_2_O_2_ release in soleus muscle. Mitochondrial oxygen consumption, membrane potentials (ΔΨ) and peroxide release were measured as described in the methodology. Blue or red bars and points represent data obtained from AL or CR animals, respectively. **A.** Oxygen consumption rates were measured in mitochondria incubated, successively, in the presence of succinate and rotenone plus ADP (State 3), oligomycin (State 4) and CCCP (Maximal). **B.** Respiratory control ratios (State 3/State 4). **C.** Membrane potentials, calibrated in mV. **D.** Hydrogen peroxide release, normalized to average AL fluorescence. ns = non-significant, ** p<0.005 versus AL.

## Discussion

We have previously found that CR increased both calcium uptake rates and maximal uptake capacity in isolated brain and liver mitochondria (Amigo *et al.*, 2017; Menezes *et al.*, 2017). Given the central role of both mitochondria and calcium ions in the regulation of energy metabolism, we find the modulation of Ca^2+^ transport in mitochondria by a dietary intervention important. As a result, we investigated if this change was observed in muscle tissues, in which rich Ca^2+^-induced metabolic changes are also observed. Surprisingly, neither tissue displayed any change in mitochondrial Ca^2+^ transport properties with the dietary intervention, although some bioenergetic changes were observed, namely a decrease in phosphorylating oxygen consumption rates and, in heart, decreased H_2_O_2_ release.

It should be noted that isolation of mitochondria from the tissue can change mitochondrial morphology and distribution in the cell (Kuznetsov *et al.* 2008), factors affecting mitochondrial Ca^2+^ uptake properties (Kowaltowski *et al.* 2019; Favaro *et al.* 2019). Thus, our experimental setup does not eliminate the possibility of *in vivo* differences. However, changes in isolated mitochondrial Ca^2+^ transport induced by CR and fasting were measured in isolated mitochondrial from brain and liver previously, showing that these can be independent of *in vivo* morphology (Amigo *et al.*, 2017; Menezes-Filho *et al.*, 2017). This suggests that changes seen in mitochondrial Ca^2+^ transport promoted by CR are indeed tissue-specific.

A prior publication (Hofer *et al.*, 2009) explored the effects of CR on mitochondrial calcium transport and mPT in the heart. This work differed from ours in the sense that the diet adopted was the NIH CR protocol, which involves supplementation with vitamins but not minerals in the CR group, which can significantly change mitochondrial metabolic properties (Cerqueira and Kowaltowski, 2010). Furthermore, the paper compared different fractions of mitochondria isolated from animals under distinct feeding states: fasted CR compared against non-fasted AL rats (Hofer *et al.*, 2009). Recent work from our group in liver indicates that substantial remodeling of mitochondrial function occurs with a few hours of fasting (Menezes-Filho *et al.*, 2019), and we do not know if these changes could also be present in heart. Nonetheless, the overall results obtained by both our group and theirs are consistent in the sense that only small or no changes in heart mitochondrial Ca^2+^ transport are promoted by CR.

In heart, mitochondrial hydrogen peroxide release rates were decreased by CR, while in soleus muscle mitochondria no such changes were detected. The available literature regarding redox balance and CR in heart mitochondria is mixed, with some works reporting decreased or even unaltered reactive oxygen species release in response to CR (Walsh *et al.* 2014). The same inconsistent results have been reported in the literature for skeletal muscle (Walsh *et al.* 2014). To to our knowledge, no prior reports were available related to the effects of CR on soleus muscle redox balance. However, our data contrast with work that measured H_2_O_2_ release in Male Fischer 344 Brown x Norway hybrid (F344×BN F1) rats (Hofer *et al.*, 2009) and observed a decrease in peroxide release promoted by CR. Notably, these changes were significant only at 18 and 29 months of age, a much more advanced time point than analyzed by us. This suggests changes in H_2_O_2_ release promoted by CR in muscle may be a consequence of different aging rates with CR, rather than an early effect of the intervention.

Both in heart and skeletal muscle, we measured moderate decreases in respiratory rates under phosphorylating conditions. Mixed results have also been reported previously for CR effects on heart mitochondrial respiration (Desai 1996; Judge *et al.*, 2004; Hofer *et al* 2009; Lanza *et al* 2012; Ruetenik *et al.* 2015). The predominant view includes non-modified or even increased respiratory rates induced by this diet, which makes our finding surprising. Many of these works, however, employed different substrates, different kinds of mitochondrial preparations, other species or strains, other forms of dietary limitation (Cerqueira and Kowaltowski, 2010), as well as other treatment durations. Indeed, bioenergetic changes induced by CR in soleus muscle were not available to date, to our knowledge.

## Conclusions

We measured mitochondrial respiration, inner membrane potentials and H_2_O_2_ in heart and soleus muscle, adding upon data in the literature and demonstrating that many tissue, substrate and diet-specific changes in bioenergetics and oxidant production exist. We also measured mitochondrial Ca^2+^ uptake, including maximal uptake capacity, Ca^2+^ uptake rates, and affinity. Interestingly, we found that these were unchanged by CR, a result that contrasts sharply with brain and liver, in which CR strongly enhances Ca^2+^ uptake in mitochondria. These results highlight the strong tissue-specificity of mitochondrial effects of CR.

## Declarations

### Funding

Supported by the Fundação de Amparo à Pesquisa do Estado de São Paulo (FAPESP) grant number 2016/18633-8, Conselho Nacional de Pesquisa e Desenvolvimento (CNPq) grant number 440436/2014, Coordenação de Aperfeiçoamento de Pessoal de Nível Superior (CAPES) finance code 001, and the Centro de Pesquisa, Inovação e Difusão de Processos Redox em Biomedicina - CEPID Redoxoma, grant 2013/07937-8. JDCS is supported by a FAPESP fellowship.

### Conflicts of interest

none

### Availability of data and material

raw data are fully available upon request

### Code availability

none

### Authors’ contributions

All authors participated in experimental design, data analysis, reading and approval of the final manuscript. JDCS and CCCS conducted experiments.

